# *Trypanosoma cruzi* senses intracellular heme levels and regulates heme homeostasis by modulating *Tc*HTE protein expression

**DOI:** 10.1101/2020.01.10.901934

**Authors:** Lucas Pagura, Evelyn Tevere, Marcelo L. Merli, Julia A. Cricco

## Abstract

Heme is an essential cofactor for many biological processes in aerobic organisms. Unlike most organisms, which can synthesize it *de novo* through a conserved pathway, the etiological agent of Chagas disease, *Trypanosoma cruzi*, as well as other trypanosomatids relevant for human health, are heme auxotrophs; thereby they must import it from the hosts. *Tc*HTE protein is involved in *T. cruzi* heme transport, although its specific role remains elusive. In the present work we studied endogenous *Tc*HTE in the different life cycle stages of the parasite in order to gain insight in its function in heme transport and homeostasis. We have confirmed that *Tc*HTE is predominantly detected in replicative stages (epimastigote and amastigote). We have also demonstrated that *T. cruzi* epimastigotes can sense intracellular heme content by an unknown mechanism and regulates heme transport to adapt to changing conditions. Based on these results, we propose a model in which *T. cruzi* senses intracellular heme and regulates heme transport activity adjusting the expression of *Tc*HTE. The elucidation and characterization of heme transport and homeostasis will contribute to a better understanding of *T. cruzi* biology as well as other trypanosomatids, pointing out this pathway as a novel drug target for therapeutics.

## Introduction

*Trypanosoma cruzi* is a protozoan parasite responsible of Chagas disease. It is a trypanosomatid, members of the class Kinetoplasteae that also includes others relevant for human health such as *Trypanosoma brucei* (sleeping sickness) and *Leishmania spp*. (visceral, cutaneous, and muco-cutaneous leishmaniasis). *T. cruzi* undergoes a complex life cycle, alternating between a mammal host and an insect vector, and displaying at least four developmental stages that are morphologically and metabolically different: epimastigotes, metacyclic trypomastigotes, intracellular amastigotes and bloodstream trypomastigotes (1). Trypanosomatids are aerobe organisms that present several heme-proteins involved in essential metabolic pathways (2). However, they lack a complete heme synthesis pathway (3); for this reason they must scavenge this molecule from the host or vector (2).

In the last years, some proteins of trypanosomatids displaying sequence homology to *Ce*HRG-4 (*<u>C</u>aenorhabditis <u>e</u>legans* <u>H</u>eme <u>R</u>esponsive <u>G</u>ene 4) (4) have been studied. LHR1 (*<u>L</u>eishmania* <u>H</u>eme <u>R</u>esponse 1, from *Leishmania amazonensis*) was the first protein postulated as a heme transporter in *Leishmania spp*. LHR1 mRNA responds to heme availability in the environment, confirming it as a member of the HRG family (5). Also, heme uptake mediated by LHR1 results relevant for the virulence of *L. amazonensis* (6, 7). We previously described an homologous protein in *T. cruzi* named *Tc*HTE (*<u>T</u>. <u>c</u>ruzi* <u>H</u>eme <u>T</u>ransport Enhancer), that plays a critical role in heme transport and is found mainly in the flagellar pocket of *T. cruzi* epimastigotes (8). More recently, *Tb*HRG (*<u>T</u>. <u>b</u>rucei* <u>H</u>eme <u>R</u>esponsive <u>G</u>ene) was described in *T. brucei* and postulated to be involved in transport of free-heme or in the salvage of heme derived from hemoglobin degradation (9, 10). LHR1, *Tc*HTE and *Tb*HRG present similar growth performance when expressed in *Saccharomyces cerevisiae hem1*Δ, allowing these knock out cells to grow in a medium supplemented with low heme. Also, the over-expression of recombinant versions of these trypanosomatid genes in the corresponding native organisms causes an increment in the intracellular heme concentration, confirming their role in heme transport (5, 8, 9). Bioinformatic studies have shown that these trypanosomatid proteins present significant similarities to the predicted topology of *Ce*HRG-4 (4), such as 4 trans-membrane domains (5, 8, 9). However, it remains unclear how the HRG proteins work, which are their precise roles in heme transport and how the expression of HRG genes is regulated by heme.

In this work, we present the study of the endogenous *Tc*HTE. Our data clearly demonstrate that *Tc*HTE mRNA directly responds to heme similar to how LHR1 does (5), confirming *Tc*HTE as a member of HRG family. Protein level also adjusts to heme transport requirements. Using different heme fluorescent analogs, we proved that epimastigotes sense intracellular heme concentration and consequently modulate the amount of *Tc*HTE. Besides, the expression of recombinant *Tc*HTE enhances replication of intracellular amastigotes, probably by increasing heme uptake from the cytoplasm of the infected cell, where its availability could be a limiting growth factor. In summary, our results show that heme transport activity in *T. cruzi* is tightly modulated by the presence or absence of *Tc*HTE.

## Results

### The accumulation of *Tc*HTE (mRNA and protein) changes through the different life-cycle stages of *Trypanosoma cruzi*

*Tc*HTE was previously described as a critical protein for heme transport in *T. cruzi*, presumably being part of the heme transporter and/or regulating its activity (8). To expand our knowledge of *Tc*HTE, we used specific polyclonal antibodies against *Tc*HTE to detect and analyze the presence of the endogenous protein along the *T. cruzi* life-cycle stages. The Western blot assays revealed that *Tc*HTE was highly expressed in amastigotes and epimastigotes (replicative stages) and almost undetectable in trypomastigotes (the infective and non-replicative stage), as it is shown in Figure 1A. *Tc*HTE expression varied together with hemin concentration in the medium when epimastigotes were incubated 3 days in LIT-10% FBS with 0, 5, and 20 μM hemin. The signal corresponding to this protein was more intense at lower hemin concentrations. Also, we analyzed the accumulation level of *Tc*HTE mRNA by qRT-PCR assay. The amount of *Tc*HTE mRNA was significantly higher in epimastigotes than amastigotes. Also, the amounts detected in epimastigotes and amastigotes were significantly higher compared to trypomastigotes, as it is shown in Figure 1B. These results indicate that *Tc*HTE was upregulated at mRNA and protein level in the replicative stages compared to the infective stage.

**Figure 1.**
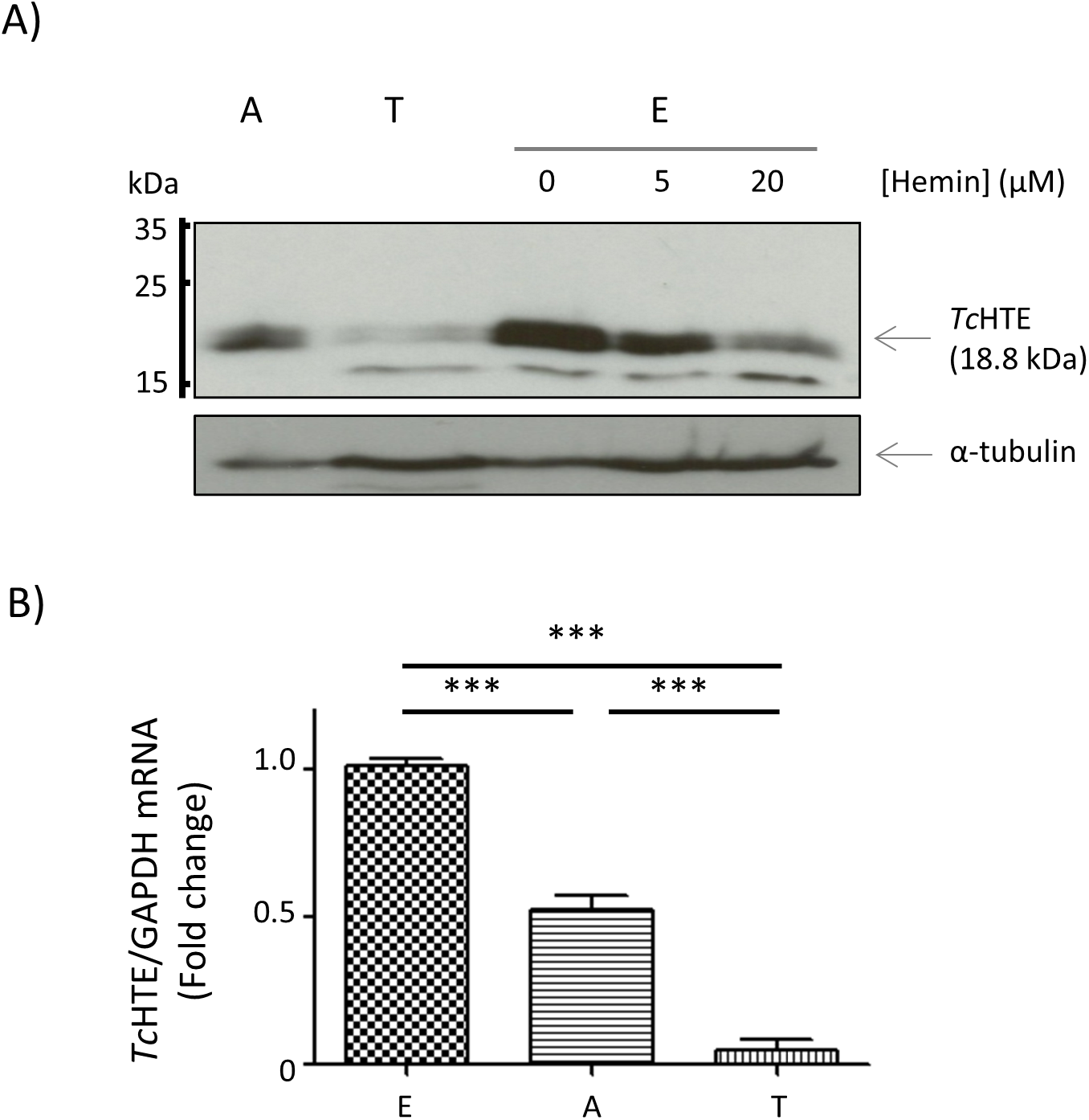
*Tc*HTE is upregulated in the replicative life-cycle stages. *A*, Western Blot analysis using anti-*Tc*HTE antibodies to detect the endogenous expression of *Tc*HTE in different life-cycle stage of *T. cruzi:* intracellular amastigotes (A), trypomastigotes (T), and epimastigotes (E). The epimastigotes were incubated in axenic culture for 3 days in LIT-10% FBS supplemented without or with 5 and 20 μM hemin (0, 5, and 20). anti-tubulin was used as a loading control. *B*, qRT-PCR analysis to detect *Tc*HTE transcript level in epimastigotes from axenic culture (E), intracellular amastigotes (A), and trypomastigotes (T) of *T. cruzi*. Transcripts from intracellular amastigotes and cells derived trypomastigotes were compared with epimastigotes grown in LIT-10% FBS plus 5 μM hemin, which was set to 1. GAPDH was used for normalization. Data is presented as mean ± SD of three independent assays. Statistical significance was determined by One-way ANOVA followed by Bonferroni’s Multiple Comparisons Test (p < 0.0001).

### Variation in the concentration of hemin affects the amount of *Tc*HTE detected in replicative life-cycle stages

To explore how changes in the amount of hemin added to the culture medium modulate the amount of *Tc*HTE present in cells, we first reviewed the growth conditions reported for epimastigotes. Previously, we have shown that *T. cruzi* incorporates heme and heme analogues (HAs) in the replicative life-cycle stages to fulfill the requirements for this cofactor. The addition of 5-20 μM hemin to the medium did not affect epimastigotes growth and, in stationary state, the intracellular heme concentration was approximately the same in both cases. However, higher concentrations of hemin produced a growth defect, being highly toxic at 100 μM (8). Then, we first analyzed the effect caused by an increment in hemin concentration in the medium but below lethal concentrations (50 μM or less). Epimastigotes routinely maintained in a medium with 5 μM hemin were collected, washed, and starved for heme for 72 h (medium without hemin). After that, parasites were cultured in media with 0, 5, 20 or 50 μM hemin for seven days. Then, cultures were diluted in fresh media (maintaining the same hemin concentration) and the growth was followed until day 14 (Figure 2A). At day 7, slight differences in the total parasite number were observed between epimastigotes growing in media with 0, 5, and 20 μM hemin and also a moderate negative effect at 50 μM hemin. However, during the second week, the negative effect produced by the absence (no hemin added) and higher hemin concentrations (20 and 50 μM) was more severe. The addition of 5 μM hemin resulted the optimal concentration to allow parasite’s growth in our laboratory conditions. Also, on the third and seventh days of the experiment, samples were collected, stained with Giemsa reagent, and analyzed by optical microscopy (Figure 2B). The epimastigotes challenged to growth in media with 0 or 5 μM hemin presented a conserved elongated shape, as typically described for this stage. However, those maintained in media with 20 or 50 μM hemin presented a rounded shape, possibly as a consequence of a potential toxic effect caused by the amount of hemin present in these media (11). *Tc*HTE protein accumulation was analyzed by Western blot assays after 72 h of incubation in the mentioned conditions (Figure 2C, left panel shows incubations scheme). The protein signal was observed in samples taken from 0 and 5 μM hemin conditions, but it was undetectable in 20 or 50 μM hemin, as it is shown in Figure 2C (right panel), in agreement with results reported in Figure 1A.

**Figure 2.**
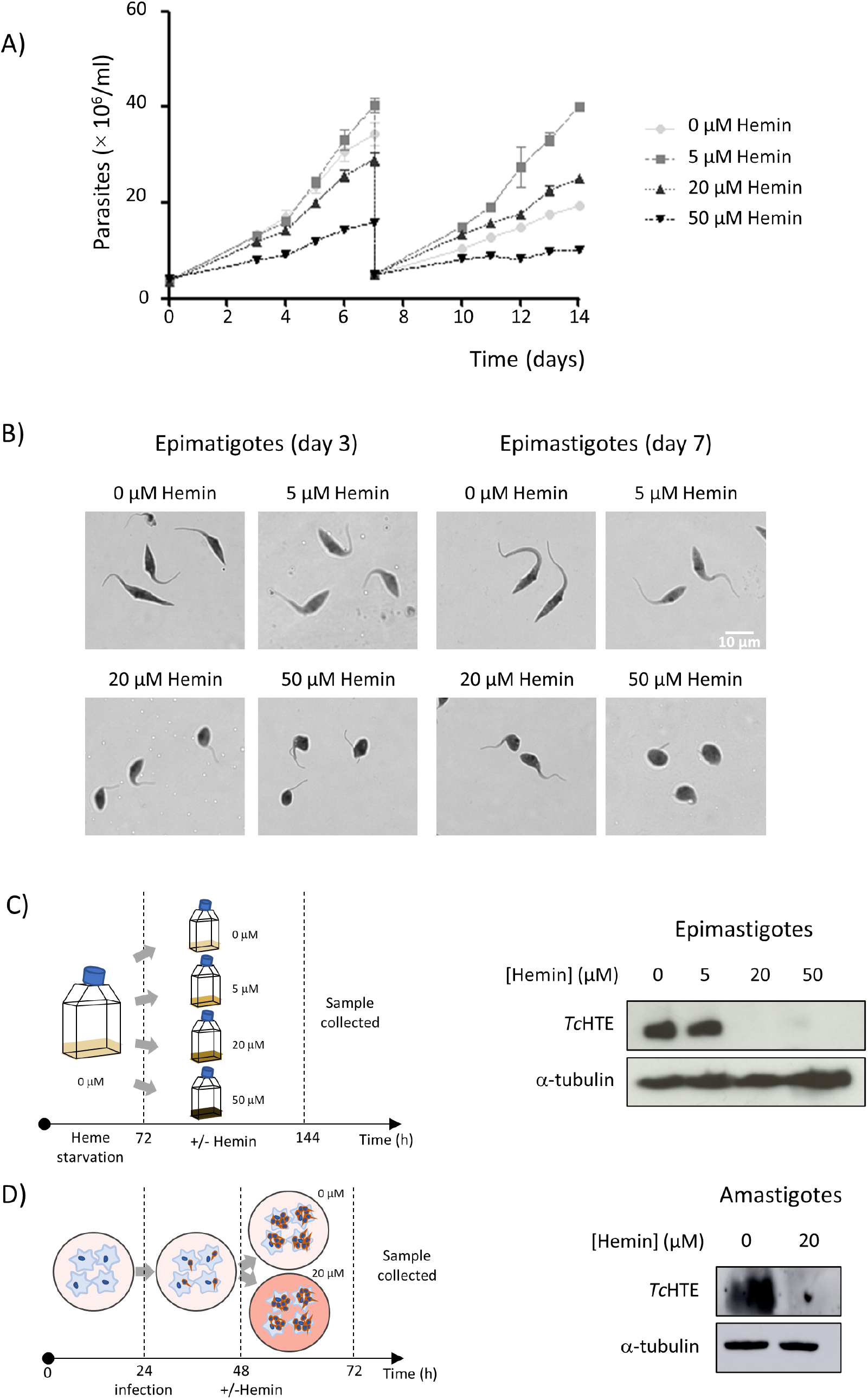
Changes in heme concentration in the medium affects epimastigote proliferation and *Tc*HTE expression. *A*, Growth curves of epimastigotes in LIT-10% FBS plus 0, 5, 20, and 50 μM hemin, with a dilution to the initial concentration in fresh medium at day 7. Previously, parasites were maintained 72 h in LIT-10% FBS without hemin. Data is presented as mean ± SD of three independent assays. *B*, Giemsa staining of epimastigotes cultured in LIT-10% FBS with 0, 5, 20, or 50 μM hemin for 3 and 7 days. *C*, Scheme of the experimental design to analyze endogenous *Tc*HTE expression in epimastigotes (left panel). Western Blot analysis (right panel) using anti-*Tc*HTE antibodies shows the accumulation of *Tc*HTE protein in total extracts of epimastigotes incubated for 72 hours without or with low hemin. anti-tubulin was used as a loading control. *D*, Scheme of the experimental design to analyze endogenous *Tc*HTE expression in amastigotes (left panel). Western Blot analysis (right panel) using anti-*Tc*HTE antibodies shows the accumulation of *Tc*HTE protein in total extracts of amastigotes incubated for 24 hours without or with 20 μM hemin. anti-tubulin was used as a loading control.

Analyzing the effect of heme on *Tc*HTE expression in intracellular amastigotes presents a difficulty because the addition of hemin to the medium during the whole assay (infection and replication) results in cell toxicity. Then, the accumulation of *Tc*HTE in this stage was evaluated in samples taken from infected cells that were treated without (as control) or with 20 μM hemin for only 24 h (Figure 2D left panel shows the scheme of treatment). The amastigotes were purified and the presence of *Tc*HTE was analyzed by Western blot. The signal corresponding to *Tc*HTE was clearly detected in the samples corresponding to amastigotes control, but the addition of 20 μM hemin caused a significant reduction in the amount of *Tc*HTE, being almost undetectable, as it is shown in the right panel of Figure 2D. In summary, *Tc*HTE was detected in the replicative forms of *T. cruzi*, and it was more abundant under heme deprivation or low heme availability.

### The amount of *Tc*HTE (mRNA and protein) in epimastigotes responds to changes in the concentration of hemin

To investigate how *Tc*HTE expression responds to hemin we analyzed if the accumulation of *Tc*HTE mRNA was affected when epimastigotes are challenged to a medium containing lower or higher concentration of hemin. Briefly, epimastigotes maintained with 0 or 20 μM hemin for 72 h were collected, washed, and incubated with 0, 5, and 20 μM hemin for 18 h. In both cases, changes in the amount of *Tc*HTE mRNA relative to GAPDH (housekeeping gene) were quantified by qRT-PCR and results are presented in Figure 3A. When epimastigotes maintained with 20 μM hemin were changed to a medium without hemin (0 μM), the amount of *Tc*HTE mRNA significantly increased, at least 3 times (Figure 3A, right panel). On the other hand, when epimastigotes maintained without hemin were changed to 5 or 20 μM hemin, the amount of accumulated *Tc*HTE mRNA significantly decreased in both cases, approximately 2 and 3 times, respectively (Figure 3A, left panel). These results confirmed *Tc*HTE as a member of HRG family since mRNA accumulation was regulated by heme in *T. cruzi* (Supplementary Information S2, phylogenetic tree of representative HRGs proteins).

**Figure 3.**
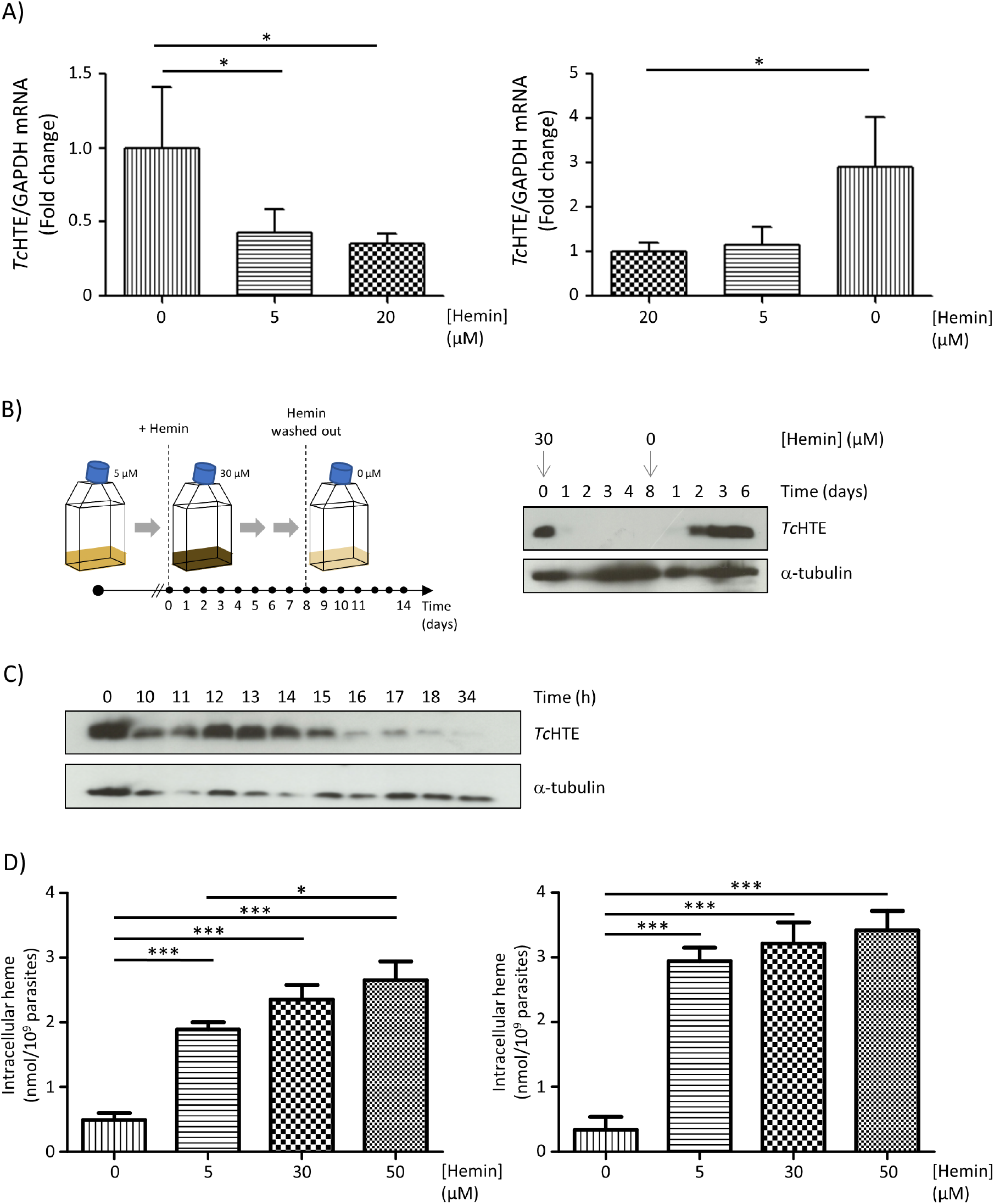
*Tc*HTE expression responds to heme at mRNA and protein level in epimastigotes. *A*, Quantification of *Tc*HTE mRNA level in epimastigote with different concentrations of hemin. Epimastigotes were cultured in LIT-10% FBS without (left panel) or with 20 μM hemin (right panel). Three days after, cells were collected, washed with PBS, and cultured in LIT-10% FBS supplemented with 0, 5 or 20 μM heme for 18 h and *Tc*HTE mRNA was quantified by qRT-PCR. GAPDH was used for normalization. Data is presented as mean ± SD of three independent assays. Statistical significance was determined by One-way ANOVA followed by Bonferroni’s multiple comparisons test (* p < 0.05). *B*, Quantification of *Tc*HTE protein level in epimastigote with different concentration of hemin. Scheme of the experimental design to analyze endogenous *Tc*HTE protein expression in epimastigote stage. Parasites were growth in LIT-10% FBS plus 5 μM hemin and then incubated in LIT-10% FBS plus 30 μM hemin (day 0). Samples were taken every day for a week. The remaining epimastigotes were washed and maintained in LIT-10% FBS without hemin (day 8) and samples were taken every day for another week (left panel). Western blot assay (right panel) using anti-*Tc*HTE antibodies to recognize endogenous protein in total extracts of epimastigotes. anti-tubulin was used as loading control. *C*, Epimastigotes grown in LIT-10% FBS supplemented with 5 μM hemin were then incubated in LIT-10% FBS plus 30 μM hemin. Samples were taken every hour. Western blot assay using anti-*Tc*HTE antibodies to recognize endogenous protein in total extracts of epimastigotes. anti-tubulin was used as loading control. *D*, Intracellular heme content determined by pyridine method in epimastigotes grown in LIT-10% FBS plus 0, 5, 30, and 50 μM hemin, for 18 h (left panel) and 24 h (right panel). Data is presented as mean ± SD of three independent assays. Statistical significance was determined by One-way ANOVA followed by Tukey’s Multiple Comparisons Test (*** p < 0.01).

Additionally, changes in *Tc*HTE protein expression were evaluated by Western blot assays. Briefly, epimastigotes routinely maintained in medium with 5 μM hemin were transferred to 30 μM hemin for 7 days. On day 8, epimastigotes were washed and transferred to fresh medium without hemin (0 μM) for 7 days. Samples were taken every day along the whole assay (left panel of Figure 3B shows the scheme of this treatment). The results, presented in Figure 3B (right panel), show that the protein signal almost disappeared 24 h after epimastigotes were incubated with 30 μM hemin. On the other hand, when epimastigotes were transferred to a hemin-free medium, 48 h were required to restore *Tc*HTE signal. To follow the drop of *Tc*HTE during the first day of increasing hemin, epimastigotes maintained with 5 μM were transferred to 30 μM hemin, samples were taken every hour and treated for Western blot assays. Figure 3C clearly shows that *Tc*HTE signal became almost undetectable 15-17 h after hemin concentration was increased.

Furthermore, we quantified intracellular heme in epimastigotes (which were previously starved for heme for 48 h) after 18 and 24 h of incubation with 0, 5, 30, and 50 μM hemin. Intracellular heme significantly increased in samples incubated with hemin (18 and 24 h) compared to the sample maintained without hemin (Figure 3D, both panels). Also, analysis of samples taken at 18 h of incubation showed a significant difference in intracellular heme of epimastigotes incubated with 5 and 50 μM hemin (Figure 3D, left panel). This difference disappeared after 24 h of incubation (Figure 3D, right panel) and intracellular heme reached almost the same concentration of approximately 3 nmol/10^9^ parasites.

### Epimastigotes can sense intracellular heme (and HAs) to modulate *Tc*HTE protein level

There is no direct evidence of any intracellular or extracellular signal that might trigger changes in *Tc*HTE accumulation. We have previously shown that fluorescent heme analogs (HAs) that have been used to evaluate heme transport activity in *T. cruzi* (8, 12, 13), can be selectively incorporated by the replicative stages of the parasite (8). Based on this potentiality, we designed a strategy to evaluate if epimastigotes sense intracellular or environmental heme (or HAs) to modulate *Tc*HTE expression. First, epimastigotes were challenged to grow under nontoxic concentration of “total metallo-prophyrins” of 20 μM or lower (total metallo-prophyrins is referred to hemin plus different HAs), and parasite proliferation and HAs internalization were analyzed. The growth profile of epimastigotes in a medium containing hemin (20 μM or 5μM) or 5 μM hemin + 15 μM HAs (20 μM of total metallo-prophyrins) is shown in Figure 4A. Epimastigotes incubated with 5 or 20 μM hemin or with 5 μM hemin plus 15 μM SnMP (Sn(IV) mesoporphyrin IX) did not show any significant difference in the growth profile. On the other hand, parasites incubated with 5 μM hemin plus 15 μM ZnMP (Zn(II) mesoporphyrin IX) were severely affected, presumably due to ZnMP toxicity. Samples were taken at the third day of the curve and the presence of HAs inside cells was analyzed by confocal microscopy and direct fluorescent measurement in a similar way as we previously reported for heme/HAS transport activity (8). Figures 4B and C show that epimastigotes incubated with SnMP (5 μM hemin plus 15 μM SnMP) did not exhibit any significant fluorescent signal detected by confocal microscopy or direct measurements of fluorescence, suggesting that it was not imported or it was rapidly exported. In any case, SnMP did not remain inside the cell. On the other hand, epimastigotes incubated with ZnMP (5 μM hemin plus 15 μM ZnMP) showed an intense fluorescence signal, detected by confocal microscopy and direct fluorescence measurements, confirming its incorporation by the cell. These observations are in agreement with previous results in which we have shown that ZnMP, but not SnMP, is internalized by *T. cruzi* (8). The effect of these HAs on *Tc*HTE accumulation was analyzed by Western blot assay in samples taken after three days of incubation with the HAs (0, 5, 20 μM hemin, 5 μM HAs or 5 μM hemin + 15 μM HAs) and the results are shown in Figure 4D. The signal corresponding to *Tc*HTE was detected in samples from 0 and 5 μM hemin, 5 μM of SnMP, 5 μM of hemin plus 15 μM of SnMP (20 μM total metallo-prophyrins) and with 5 μM of ZnMP conditions. On the other hand, *Tc*HTE was not detected neither in samples from 5 μM of hemin plus 15 μM of ZnMP nor 20 μM of hemin conditions. These results indicate that only those HAs that were internalized and remained in the parasite provoked a reduction on *Tc*HTE protein level similar to the reduction produced by higher hemin.

**Figure 4.**
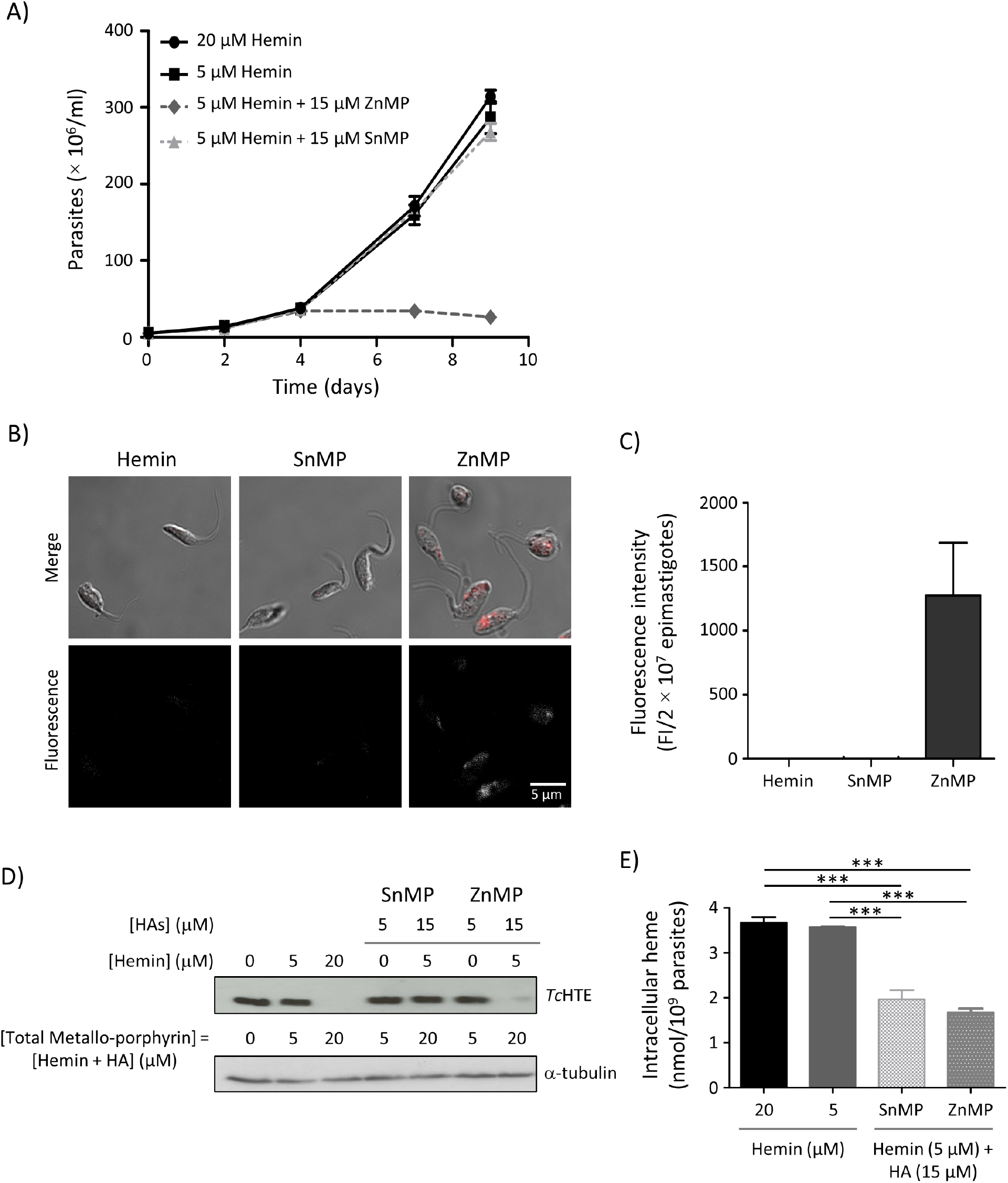
Heme analogs affect epimastigote growth and *Tc*HTE accumulation. *A*, Growth curves of *T. cruzi* epimastigotes in LIT-10% FBS plus 5, 20 μM hemin or 5 μM hemin plus 15 μM of HAs (ZnMP and SnMP), monitored for 9 days. Parasites were maintained in exponential growth by periodic dilution every two days in fresh medium. Previously, parasites were maintained 72 h in LIT-10% FBS without hemin. Data is presented as mean ± SD of three independent assays. *B*, Confocal microscopy of epimastigotes incubated with 5 μM hemin or 5 μM hemin plus 15 μM of the different HAs for 3 days. *C*, Fluorescence intensity (FI) from cell-free extracts were obtained after 3 days of incubation of epimastigotes with 5 μM hemin or 5 μM hemin plus 15 μM of the different HAs. The experimental data is presented as the mean ± SD of three independent replicates. *D*, Western blot assay, using anti-*Tc*HTE antibodies to recognize endogenous protein in total extracts of epimastigotes cultured with 0, 5 or 20 μM hemin, 5 μM Has, or 5 μM hemin plus 15 μM HAs for four days. anti-tubulin was used as a loading control. *E*, Intracellular heme content determined by pyridine method of epimastigotes grown in LIT-10% FBS plus 5, 20 μM hemin, or 5 μM hemin plus 15 μM HAs, for 72 h. Data is presented as mean ± SD of three independent assays. Statistical significance was determined by One-way ANOVA followed by Tukey’s Multiple Comparisons Test (*** p ≤ 0.01).

When we analyzed intracellular heme, we observed that the treatment with both HAs (ZnMP and SnMP) significantly reduced intracellular heme content, indicating that somehow SnMP blocked heme uptake, causing a drop in intracellular heme levels, as it is shown in Figure 4E.

### r*Tc*HTE.His-GFP enhances replication of amastigotes

The replication of intracellular amastigotes was analyzed in wild type and recombinant parasites over-expressing r*Tc*HTE.His-GFP. Briefly, monolayers of Vero cells were infected with wild type or r*Tc*HTE.His-GFP overexpressing trypomastigotes. 48 h post infection, cells were fixed and stained with Giemsa and amastigotes were counted (Figure 5A). The number of intracellular amastigotes was significantly higher in cells infected with r*Tc*HTE.His-GFP overexpressing parasites compared to those infected with wild type parasites. (29 ± 2 and 18 ±1 amastigotes/infected-cell, respectively, p<0.0001). This result suggests that the over-expression of *Tc*HTE increased amastigote replication rate in a low free-heme environment as the cellular cytosol.

**Figure 5.**
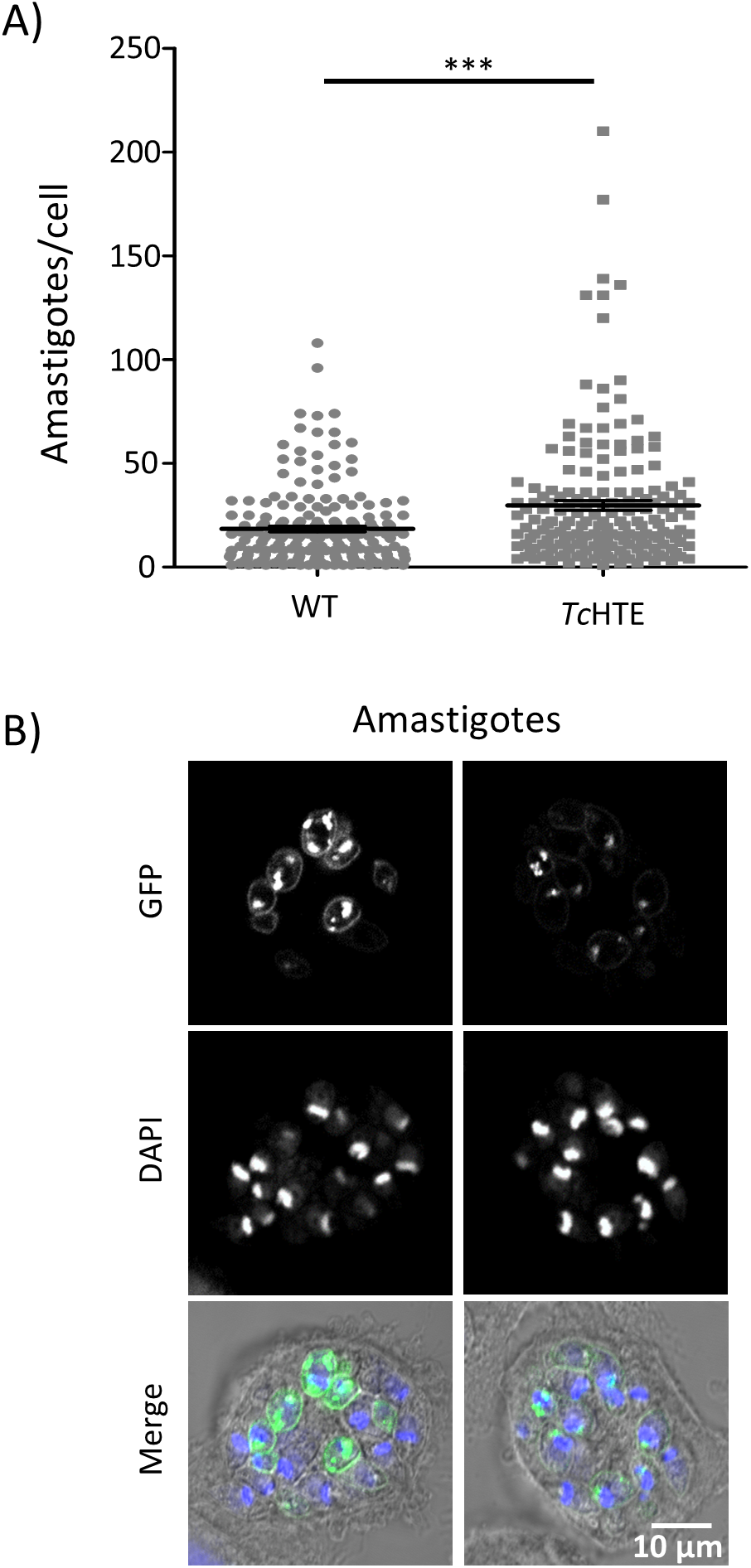
Over-expression of r*Tc*HTE.His-GFP in *T. cruzi* stimulates amastigotes proliferation. *A*, Vero cells were infected with wild type parasites or parasites that over-express r*Tc*HTE.HIS-GFP. 72 h after infection, cells were fixed and stained with Giemsa reagent. Intracellular amastigotes were counted from more than 180 cells for each condition. Statistical significance was determined by Mann-Whitney’s Test. The experimental data is presented as the mean ± SD (*** p < 0.0001). *B*, Confocal microscopy images of amastigotes expressing r*Tc*HTE.HIS-GFP. r*Tc*HTE.HIS-GFP (green) and DAPI (blue).

Additionally, we also analyzed the presence of r*Tc*HTE.His-GFP in intracellular amastigotes. The images recorded for intracellular amastigotes confirmed r*Tc*HTE.His-GFP expression and showed that its signal is highly intense and punctuated in a region over the plasma membrane that could overlap with the flagellar pocket region (Figure 5B), as it was previously shown in epimastigotes (8). However, several amastigotes also showed a fluorescent signal distributed along the plasma membrane. Unfortunately, endogenous *Tc*HTE could not be detected by indirect immunofluorescence assays, using anti-*Tc*HTE antibodies. Then, the difference observed in the localization of r*Tc*HTE.His-GFP protein in amastigotes could be an artefact of protein over-expression (r*Tc*HTE.His-GFP) or could indicate that *Tc*HTE changes its localization in the intracellular life-cycle stage.

## Discussion

Heme is an essential cofactor for all aerobic organisms and most of them are able to synthesize it *via* a conserved pathway (14–16). Also, it is well established that free-heme is not found in any living cell due to its toxicity, therefore a fine tune control of heme homeostasis is necessary to avoid its harmful effects. *T. cruzi* does not synthesize heme, it needs to take this cofactor from its mammal host or vector insect (2, 17). Once heme is inside the parasite, transporters, chaperons and/or carriers must be involved in its transport and trafficking. During the last few years, several proteins involved in heme transport have been described in trypanosomatids, such as LHR1 in *Leishmania spp*. (5–7), *Tb*HRG in *T. brucei* (9, 10), and *Tc*HTE in *T. cruzi* (8). And most recently, the protein *Lm*FLVCRb, member of the Major Facilitator Superfamily, was described as a heme importer in *L. major* (18). The latest might be involved in free-heme transport in *L. major*, probably overlapping some functions with *Lm*HR1. However, the mechanism of heme transport, the precise identity of the transporters and their regulation have not been completely unraveled neither in *T. cruzi* nor in other trypanosomatids.

In this work we present data that clarifies the relationship between *Tc*HTE and heme transport activity in *T. cruzi*. The attainment of polyclonal antibodies against *Tc*HTE allowed the evaluation of the endogenous protein. It was clearly detected in epimastigotes and amastigotes, and almost undetectable in trypomastigotes. In accordance, quantitative RT-PCR analysis showed that *Tc*HTE mRNA was significantly higher in the replicative life-cycle stages of *T. cruzi*. *Tc*HTE (as mRNA and protein) was more abundant in the stages where *T. cruzi* is able to import heme (or heme analogs HAs) (8). In addition, the amount of hemin added to the medium inversely affected the amount of *Tc*HTE mRNA detected in epimastigotes, as it was reported for LHR1 promastigotes (5). These results confirmed *Tc*HTE as a member of the <u>H</u>eme <u>R</u>esponse <u>G</u>ene (HRG) family (4). It is important to note that *Leishmania* (5) and *T. cruzi* HRGs respond to heme but not *Tb*HRG (10), suggesting that those gene products might be under different regulations and/or play different roles in heme transport.

Epimastigotes incubated with different concentrations of hemin reached similar intracellular heme concentration. This suggests that *T. cruzi* might exhibit an optimal intracellular heme concentration or “heme quota” despite the amount of heme available in the medium. Under heme starvation treatment, intracellular heme drops below the optimal, triggering the expression of *Tc*HTE (allowing its detection by Western blot). Once the internal heme quota is satisfied, *Tc*HTE protein level decreases (almost undetectable by Western blot). These results strongly suggest that heme requirement could modulate *Tc*HTE protein level (and mRNA) in the parasite, supporting a direct relation between *Tc*HTE and heme transport activity. We cannot exclude heme degradation as part of a mechanism to control intracellular heme, but genes encoding enzymes with heme oxygenase homology have not been identified in the genome of *T. cruzi* (19). Presumably, heme degradation *via* heme oxygenase like activity previously reported (19) would not be enough to get rid of any excess of heme when epimastigotes are exposed at high concentration of hemin. Under this premise, the amount of intracellular heme should be mainly controlled by its transport, with *Tc*HTE playing a role as heme transporter or as an essential part of it.

To clarify if *Tc*HTE responds to signals of extracellular or intracellular heme, we took advantage that *T. cruzi* can selectively import HAs (ZnMP, but not SnMP) (8). Interestingly, although treatment with both HA reduce intracellular heme levels, only ZnMP reduced *Tc*HTE expression. These results indicate that epimastigotes internalize ZnMP and hemin indistinctly, contributing to the concentration of “total metallo-porphyrins” (heme + ZnMP) and both are sensed by the parasite. On the other hand, SnMP is not detected inside the cell, and has no effect on *Tc*HTE expression suggesting that it is not been sensed. The fact that SnMP is not internalized or maintained inside the cell is also consistent with the fact that the growth profile is not affected. These results strongly suggest that epimastigotes only sense intracellular heme, independently of the amount of hemin present in the medium, and modulate *Tc*HTE expression to adjust heme transport activity. These results reinforce the essential role of *Tc*HTE controlling heme homeostasis in *T. cruzi*.

The treatment of infected cells with hemin caused a drop in the *Tc*HTE signal in intracellular amastigotes. Assuming that *Tc*HTE mediates heme transport activity in this life stage, the regulation of *Tc*HTE in amastigotes resembles that observed in epimastigotes. Additionally, a significantly higher number of intracellular amastigotes was determined in cell lines infected with the recombinant parasites compared with those infected with WT parasites. Since the presence of recombinant *Tc*HTE increased heme uptake in epimastigotes (8), we hypothesized that r*Tc*HTE.His-GFP also enhances heme uptake in amastigotes, therefore stimulating cellular replication. A similar evidence was found in *L. amazonensis*, in which the deletion of one copy of genomic LHR1 impaired intracellular replication of *L. amazonensis* amastigotes (7). In summary, it is reasonable to postulate that the ability of intracellular amastigotes to incorporate more heme enhances its replication rate.

Based on the results presented here, we propose a model for the role of *Tc*HTE in *T. cruzi* heme transport that is represented in Figure 6. In epimastigotes under stationary state of heme flux, protein levels of *Tc*HTE remain low, almost undetectable by Western blot. The parasite maintains a constant intracellular heme concentration with low heme import activity. Under heme deprivation (heme starvation) epimastigotes sense low intracellular heme, the expression of *Tc*HTE increases, more protein is assembled in the flagellar pocket region (as heme transporter or part of it) and heme transport activity raises. Once the intracellular heme quota is satisfied, an intracellular signal (still unknown) triggers the mRNA and also the extra-protein degradation and heme transport activity decreases to stationary level. Based on its size and its predicted structural features (171 amino acids, 4 predicted transmembrane domains (8)), *Tc*HTE should assemble as a homo-trimer to form a channel or pore of the heme transporter. Also, *Tc*HTE could be part of a hetero-complex heme transporter. Nevertheless, we cannot exclude *Tc*HTE acting only as a regulatory subunit of the transporter. In this scenario, when heme transport activity is required, *Tc*HTE is synthesized and assembled to build up or activate the transporter. The constant detection of endogenous *Tc*HTE in intracellular amastigotes is consistent with the fact that available heme in the cytoplasm of host cells is very low (labile-heme was estimated in different eukaryotic cells, in yeast 20 - 40 nM (20), HEK-293 cells approximately 450 nM (21), IMR90 lung fibroblast cells, approximately 614 nM (22)) and its presence is necessary to guarantee enough heme uptake in this life-cycle stage. The reduction of this protein signal when extra hemin was added to the infected cells supports our model for *Tc*HTE.

**Figure 6.**
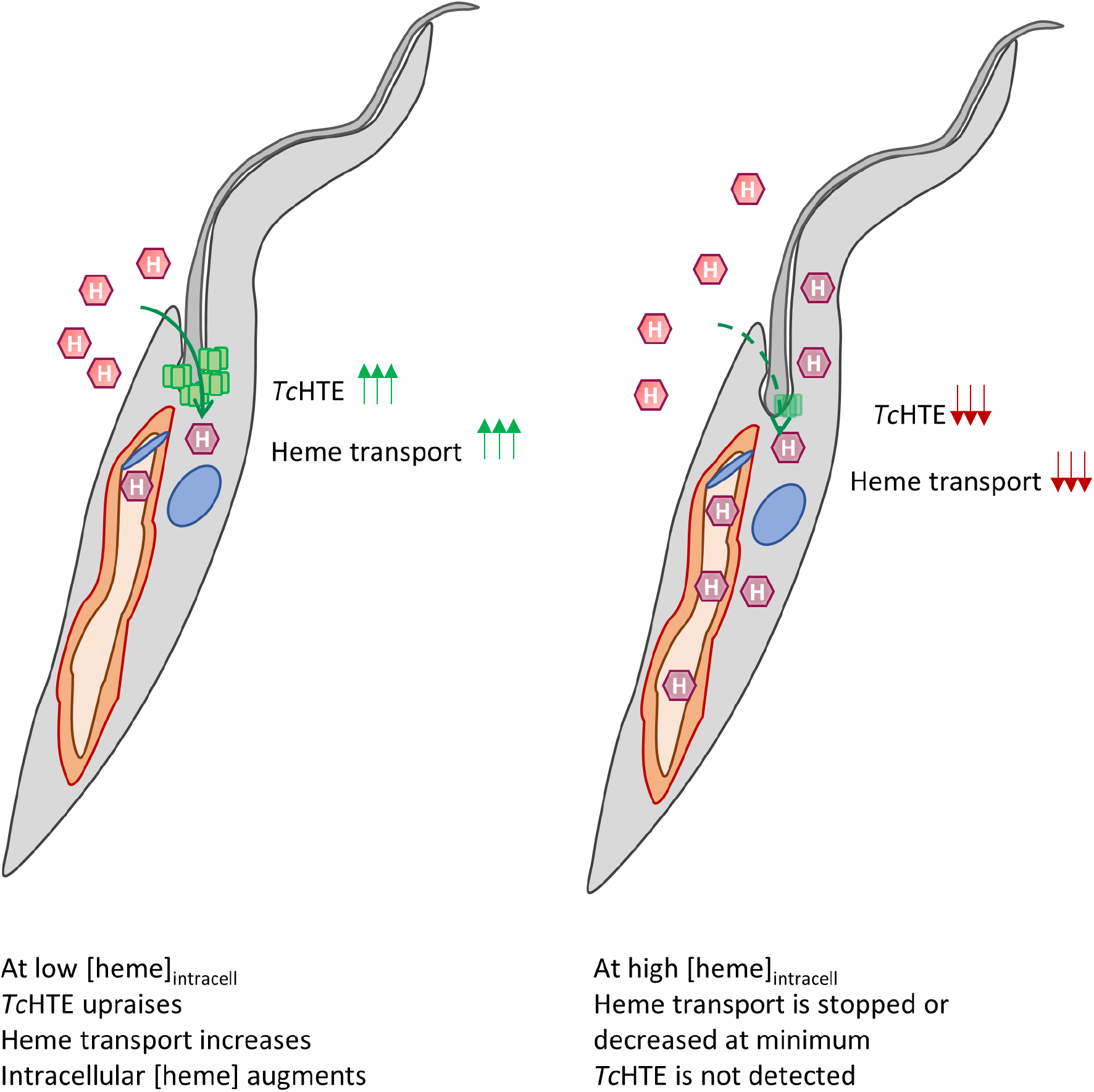
Scheme representing a proposed model for *Tc*HTE/*Tc*HRG function in heme transport. When epimastigotes sense low intracellular heme (left panel), augment *Tc*HTE synthesis and assembly in the flagellar pocket and heme uptake increases. Once intracellular heme raises the optimal quota (right panel), *Tc*HTE is downregulated and heme import is maintained low, keeping the heme flux at stationary level.

In summary, *Tc*HTE responds to intracellular heme, confirming it as a member of HRG family protein. Therefore, it might be renamed as *Tc*HRG. *T. cruzi* epimastigotes can manage intracellular heme concentration by controlling its transport in replicative life-cycle stages in which the presence of *Tc*HTE/*Tc*HRG is essential to this process. *T. cruzi* can sense intracellular heme, by a still unknown mechanism, and adjust *Tc*HTE expression and accumulation in order to promote or reduce heme transport activity. Evidences presented here strongly suggest that *Tc*HTE/*Tc*HRG can form the heme transporter or a relevant part it. In this scenario, the characterization of heme transport and distribution will contribute to a better understanding of the biology and biochemistry of *T. cruzi*, the etiological agent of the most prevalent parasitic disease in several countries of America (https://www.who.int/news-room/fact-sheets/detail/chagas-disease-(american-trypanosomiasis). This, in turn, will enable the identification of novel proteins playing essential roles which might act as possible targets for new drug development against Chagas disease.

## Experimental procedures

### Reagents

Dulbecco’s Modified Eagle Medium (DMEM) was obtained from Life Technologies, Fetal Bovine Serum (FBS) from Internegocios SA. FBS was heat-inactivated at 56°C for an hour. Hemin, ZnMP (Zn(II) mesoporphyrin IX) and SnMP (Sn(IV) mesoporphyrin IX) were obtained from Frontier Scientific. Hemin, ZnMP and SnMP stock and working solutions were prepared as previously described in (8). Heme concentration in hemin stock solution was confirmed by spectroscopic measurements at 385 nm, ε^385^= 58400 M^-1^cm^-1^ (20).

### Parasites and cell lines

All experiments were performed using *T. cruzi* Dm28c strain and all the infections were carried out in Vero cell line (ATCC CCL-81, already available in our laboratory). Epimastigotes were maintained in mid-log phase by periodic dilutions in Liver Infusion Tryptose (LIT) medium supplemented with 10% heat inactivated FBS (LIT-10% FBS) and 5 μM hemin (8), at 28 °C. The Vero cell line was routinely maintained in DMEM medium supplemented with 0.15% (w/v) NaHCO_3_,-and 10% heat inactivated FBS (DMEM-10% FBS) at 37 °C in a humid atmosphere containing 5% CO2. During the *T. cruzi* infections, Vero cells were incubated in DMEM medium supplemented 2% heat inactivated FBS (DMEM-2% FBS) at 37 °C in a humid atmosphere containing 5% CO2. Metacyclic trypomastigotes (WT and recombinant overexpressing r*Tc*HTE.HIS-GFP) were obtained by spontaneous differentiation of epimastigotes at 28 °C after two weeks of culture without dilution. Metacyclic trypomastigotes were used to infect monolayered Vero cells and to obtain cell-derived trypomastigotes after 5-6 days post-infection. Intracellular amastigotes were obtained 2- or 3-days post infection of monolayered Vero cell line with cell-derived trypomastigotes. The homogeneity of the obtained *T. cruzi* forms was evaluated by microscopic observation.

### Antibodies

Rabbit polyclonal anti-*Tc*HTE antibodies were obtained using the p1*Tc*HTE peptide (amino acid 120 to 171) expressed as a TRX.HIS-fusion protein in the vector pET-32a vector (Novagen^®^), following same strategy we described previously (23). Primers used to amplify p1*Tc*HTE are: FP 5’-TCCGGATCCATGATGGCCGCAAAGTGGTG-3’ and RP 5’-CCGCTCGAGTTAATGATGATGATGATGATGACCCATAATATCTGCGTTCTTTTCG-3’ The specificity of the antibodies were tested as shown in Supporting Figure S1. All experiments were approved by the Institutional Committee of Animal Care and Use of the Facultad de Ciencias Bioquímicas y Farmacéuticas (School of Biochemical and Pharmaceutical Science), Universidad Nacional de Rosario, Argentina, and conducted according to specifications of the US National Institutes of Health guidelines for the care and use of laboratory animals (file number 935/2015).

### Effect of heme on epimastigote growth and TcHTE protein accumulation

Epimastigotes (WT) grown in LIT-10% FBS with 5 μM hemin for 48 h were collected, washed with PBS, and challenged to grow in LIT-10% FBS without (0 μM) or supplemented with 5, 20 or 50 μM hemin. The number of cells was monitored daily for 14 days. One dilution to the initial condition was made on day 7. On the third- and seventh-days samples were collected by centrifugation at 2000 *g* for 5 minutes, washed twice with PBS and prepared for optic microscopy as described below. Also, aliquots were lysed (1 × 10^6^ parasites/μl) with lysis buffer (8 M Urea, 30 mM HEPES, pH 8) for Western Blot assays.

### Effect of heme on TcHTE protein accumulation in amastigotes

To obtain the intracellular amastigotes, Vero cells were cultured as described above. After 24 h of incubation in DMEM-2% FBS, cells were incubated with trypomastigotes WT in a ratio of 10 trypomastigotes/cell (MOI = 10, multiplicity of infection) for 4 h to let the infection progress. After that, the cells were washed twice with PBS and 2 ml of DMEM-2% FBS was added. 24 h postinfection, hemin was added at a final concentration of 20 μM. After 24 h of incubation (with or without extra hemin), cells were washed, intracellular amastigotes and protein extracts were obtained as previously described (23).

### Effect of TcHTE over-expression on amastigote proliferation

To analyze amastigote proliferation, the Vero cell line was plated on 24 well plates with coverslips in DMEM-2% FBS. After 24 h, cells were incubated 4 h with Cell-derived trypomastigotes, WT or r*Tc*HTE-over-expressing ones, at MOI = 2. After the infection, the cells were washed twice with PBS and 2 mL of fresh medium were added. 72 h postinfection, infected cells were washed with PBS, and prepared for Giemsa staining and microscopic analysis as described below. Amastigotes from 183 infected cells for each group were counted by direct observation under microscope.

### Western blot

Total protein from cell-free extracts obtained from epimastigotes, trypomastigotes and amastigotes were prepared and processed as previously described (23) with minor modifications. Samples were heated at 42 °C for 30 minutes in loading buffer and 5 × 10^6^ cells/well were resolved by electrophoresis on a 15% SDS polyacrylamide gel. *Tc*HTE detection was performed with rabbit anti-*Tc*HTE antibodies (1:5000). Bound antibodies were detected with peroxidase-labeled anti-rabbit IgG (1:30000) (Calbiochem), and ECL Prime Western Blotting Detection kit (GE Healthcare). Loading control was performed with anti-tubulin clone TAT-1 antibodies (a gift from K. Gull, University of Oxford, U.K.).

### mRNA isolation, reverse transcription PCR (RT-PCR) and quantitative real-time PCR (qRT-PCR)

The following samples were used to obtain mRNA from different life-cycle stages of *T. cruzi*: epimastigotes cultured in LIT-10% FBS supplemented with or without 20 μM hemin for 3 days were collected, washed with PBS and cultured in LIT-10% FBS supplemented with 0, 5 or 20 μM hemin for 18 h. Also, trypomastigotes, amastigotes and epimastigotes maintained in LIT-10% FBS supplemented with 5 μM hemin were used for mRNA purification as we previously described with minor modifications (24). Briefly, the total mRNA was obtained using TRI Reagent^®^ (Molecular Research Center, Inc., # TR 118). The RNA preparations were treated with RNase-free Dnase I (Fermentas, Life Sciences) and their quality and concentration were checked following standard procedures (25). Each RNA extraction was carried out by triplicate. cDNAs of *T. cruzi* were synthesized through a RT reaction (M-MuLV, Thermo-Scientific) using 0.5 μg of total RNA. mRNA analysis by quantitative Real-time PCR was performed in an Applied Biosystems StepOne™ Real-Time PCR System Thermal Cycling Block using the SYBRgreen fluorescence quantification system (Fermentas). The standard PCR conditions were: 95 °C (10 min), and then 40 cycles of 94 °C (1 min), 60 °C (1 min) and 72 °C (2 min), followed by the denaturation curve. The primer designs were based on nucleotide sequences of *T. cruzi* CL Brenner Esmeraldo-like genes coding for *Tc*HTE and glyceraldehyde-3-phosphate dehydrogenase (GAPDH) (TriTrypDB accession numbers for *Tc*HTE: TcCLB.511071.190 and GAPDH: TcCLB.506943.50). The sequences of the primers used are listed below. The data were analyzed using StepOne™ software v 2.1. The fold-change in the expression of the transcripts was obtained using the comparative method (ΔΔC_t_) (26). The epimastigote stage at 5 μM hemin was used as the reference condition for both genes. Primers for qRT-PCR: *Tc*HTEss 5’-TAATTATTGGGCGGCGGCT-3’, *Tc*HTEas 5’-GAAGTACGAACTCCCCGTCC-3’ (product 235 bp); and GAPDHss 5’-GTGGCAGCACCGGTAACG-3’, GAPDHas 5’-CAGGTCTTTCTTTTGCGAATAGG-3’ (product 110 bp). The differences in the transcriptional level among the different stages were compared using One-way Anova, followed by Bonferroni’s multiple comparison test. For this purpose, the software GRAPHPAD PRISM version 5.00 for Windows (GraphPad Software, San Diego, CA) was used. The significance level (P value) was determined with a confidence interval of 95% in a two-tailed distribution.

### Optic microscopy

Epimastigotes were fixed with formaldehyde 3.7% (w/v) in PBS, washed twice with PBS and settled on poly-L-lysine-coated microscope slides. Parasites were stained with Giemsa reagent for 20-30 minutes, washed with stabilized water and mounted for microscopic analysis. Infected cells bearing *T. cruzi* amastigotes grown over these coverslips were washed with PBS and fixed with pure methanol for 3 minutes. Then, the cells were washed again, stained with Giemsa reagent, and mounted with Canada balsam on microscope slides for microscopic analysis

### Confocal microscopy

Samples of epimastigotes and amastigotes were prepared as described previously (8) with minor modifications: infected cells with *T. cruzi* amastigotes grown over coverslips were fixed with pure methanol for 3 minutes. All the images were acquired with confocal microscopes Nikon Eclipse TE-2000-E2 or Zeiss LSM880 and were processed using the ImageJ software (27).

### Epimastigote growth curve with HAs

Epimastigotes grown in LIT-10% FBS with 5 μM hemin for 48 h were collected, washed with PBS and challenged to growth in LIT-10% FBS supplemented with 5 and 20 μM hemin, or 5 μM hemin plus 15 μM heme analogs (ZnMP and SnMP). Parasites were maintained in mid-log phase by successive dilution every 2 days and growth was monitored for 9 days. On the third day samples were collected and prepared for confocal analysis as described above or to measure direct fluorescence as described below.

### Heme content analysis

The heme content was quantified by the alkaline pyridine method described in Berry *et al*. (28) with the modification we introduced to perform this assay in epimastigotes (8). Each sample contained 150 × 10^6^ epimastigotes. As previously demonstrated, presence of HAs does not interfere with heme quantification (13).

### Fluorescence Intensity analysis

20 × 10^6^ epimastigotes treated with HAs (obtained from the growth curve) were treated and analyzed as we previously described (8). Fluorescence intensity (FI) was measure in a Varian Eclipse fluorometer: λex = 405 nm, recording the emission spectra between 450 to 650 nm and analyzing the maximal emission at 578 nm for ZnMP, and 574 nm for SnMP.

### Statistical analysis

The experiments were performed in triplicate, and the data is presented as the mean ± SD (standard deviation) except when is informed. All the assays were independently reproduced at least 2–4 times. Statistically significant differences between groups were assessed using One way ANOVA followed by Tukey’s Multiple Comparison Test (intracellular heme), Mann-Whitney’s test (amastigote proliferation) or One-way ANOVA followed by Bonferroni’s multiple comparison test (mRNA analysis), as appropriate (GRAPHPAD PRISM version 5.00 for Windows, GraphPad Software, San Diego, CA).

## Data Availability

All data are contained within the article.

## Acknowledgments

We are grateful to Rodrigo Vena for his help in the acquisition of confocal microscopy images, and also to Dolores Campos for the assistance in cell culture procedures.

## FOOTNOTES

The research leading to these results has, in part, received funding from UK Research and Innovation via the Global Challenges Research Fund under grant agreement ‘A Global Network for Neglected Tropical Diseases’ grant number MR/P027989/1, and ANPCyT (National Agency of Scientific and Technological Promotion) grant no. PICT 2015-2437 to J.A.C.

JAC is a researcher of CONICET (National Research Council of Science and Technology), LP and MLM had a fellowship from CONICET to conduct their Ph.D., ET has a fellowship of ANPCyT associated to grant no. PICT 2015-2437.

## The abbreviations used are

*Tc*HTE: *T. cruzi* Heme Transport Enhancer
r*Tc*HTE.His-GFP: recombinant *Tc*HTE
HRG: Heme-response Gene
LHR1: *Leishmania* heme response 1
*Tb*HRG: *T. brucei* heme-responsive gene
TMD: transmembrane domain
qRT-PCR: quantitative real-time PCR
SnMP: Sn(IV) mesoporphyrin
ZnMP: Zn(II) mesoporphyrin
LIT: Liver Infusion Tryptose
DMEM: Dulbecco’s modified Eagle’s medium
FBS: fetal bovine serum
DAPI: 4’,6-Diamidino-2-Phenylindole
GFP: green fluorescent protein
MOI: multiplicity of Infection
WT: wild-type

## References

1. Tyler, K. M., and Engman, D. M. (2001) The life cycle of Trypanosoma cruzi revisited. Int. J. Parasitol. 31, 472–481

2. Tripodi, K. E. J., Menendez Bravo, S. M., and Cricco, J. A. (2011) Role of heme and heme-proteins in trypanosomatid essential metabolic pathways. Enzyme Res. 2011, 873230

3. Kořený, L., Oborník, M., and Lukeš, J. (2013) Make It, Take It, or Leave It: Heme Metabolism of Parasites. PLoS Pathog. 10.1371/journal.ppat.1003088

4. Rajagopal, A., Rao, A. U., Amigo, J., Tian, M., Upadhyay, S. K., Hall, C., Uhm, S., Mathew, M. K., Fleming, M. D., Paw, B. H., Krause, M., and Hamza, I. (2008) Haem homeostasis is regulated by the conserved and concerted functions of HRG-1 proteins. Nature. 453, 1127–1131

5. Huynh, C., Yuan, X., Miguel, D. C., Renberg, R. L., Protchenko, O., Philpott, C. C., Hamza, I., and Andrews, N. W. (2012) Heme uptake by Leishmania amazonensis is mediated by the transmembrane protein LHR1. PLoS Pathog. 8, 36

6. Renberg, R. L., Yuan, X., Samuel, T. K., Miguel, D. C., Hamza, I., Andrews, N. W., and Flannery, A. R. (2015) The Heme Transport Capacity of LHR1 Determines the Extent of Virulence in Leishmania amazonensis. PLoS Negl. Trop. Dis. 9, e0003804

7. Miguel, D. C., Flannery, A. R., Mittra, B., and Andrews, N. W. (2013) Heme uptake mediated by lhr1 is essential for leishmania amazonensis virulence. Infect. Immun. 81, 3620–3626

8. Merli, M. L., Pagura, L., Hernández, J., Barisón, M. J., Pral, M. F., Silber, A. M., and Cricco, J. A. (2016) The Trypanosoma cruzi Protein TcHTE Is Critical for Heme Uptake. PLoS Negl. Trop. Dis. 10.1371/journal.pntd.0004359

9. Cabello-Donayre, M., Malagarie-Cazenave, S., Campos-Salinas, J., Gálvez, F. J., Rodríguez-Martínez, A., Pineda-Molina, E., Orrego, L. M., Martínez-García, M., Sánchez-Cañete, M. P., Estévez, A. M., and Pérez-Victoria, J. M. (2016) Trypanosomatid parasites rescue heme from endocytosed hemoglobin through lysosomal HRG transporters. Mol. Microbiol. 101, 895–908

10. Horáková, E., Changmai, P., Vancová, M., Sobotka, R., Abbeele, J. Van Den, Vanhollebeke, B., and Lukeš, J. (2017) The Trypanosoma brucei Tb Hrg protein is a heme transporter involved in the regulation of stage-specific morphological. J. Biol. Chem. 292, 6998–7010

11. Schmitt, T. H., Frezzatti, W. A., and Schreier, S. (1993) Hemin-Induced Lipid Membrane Disorder and Increased Permeability: A Molecular Model for the Mechanism of Cell Lysis. Arch. Biochem. Biophys. 307, 96–103

12. Lara, F. A., Sant’Anna, C., Lemos, D., Laranja, G. A. T., Coelho, M. G. P., Reis Salles, I., Michel, A., Oliveira, P. L., Cunha-e-Silva, N., Salmon, D., and Paes, M. C. (2007) Heme requirement and intracellular trafficking in Trypanosoma cruzi epimastigotes. Biochem. Biophys. Res. Commun. 355, 16–22

13. Cupello, M. P., Souza, C. F. De, Buchensky, C., Soares, J. B. R. C., Laranja, G. A. T., Coelho, M. G. P., Cricco, J. A., and Paes, M. C. (2011) The heme uptake process in Trypanosoma cruzi epimastigotes is inhibited by heme analogues and by inhibitors of ABC transporters. Acta Trop. 120, 211–218

14. Hamza, I., and Dailey, H. a. (2012) One ring to rule them all: Trafficking of heme and heme synthesis intermediates in the metazoans. Biochim. Biophys. Acta - Mol. Cell Res. 1823, 1617–1632

15. Choby, J. E., and Skaar, E. P. (2016) Heme Synthesis and Acquisition in Bacterial Pathogens. J. Mol. Biol. 10.1016/j.jmb.2016.03.018

16. Panek, H., and O’Brian, M. R. (2002) A whole genome view of prokaryotic haem biosynthesis. Microbiology. 148, 2273–2282

17. Kořený, L., Lukeš, J., and Oborník, M. (2010) Evolution of the haem synthetic pathway in kinetoplastid flagellates: An essential pathway that is not essential after all? Int. J. Parasitol. 40, 149–156

18. Cabello-Donayre, M., Orrego, L. M., Herráez, E., Vargas, P., Martínez-García, M., Campos-Salinas, J., Pérez-Victoria, I., Vicente, B., Marín, J. J. G., and Pérez-Victoria, J. M. (2019) Leishmania heme uptake involves LmFLVCRb, a novel porphyrin transporter essential for the parasite. Cell. Mol. Life Sci. 10.1007/s00018-019-03258-3

19. Cupello, M. P., Souza, C. F., Menna-Barreto, R. F., Nogueira, N. P. A., Laranja, G. A. T., Sabino, K. C. C., Coelho, M. G. P., Oliveira, M. M., and Paes, M. C. (2014) Trypanosomatid essential metabolic pathway: New approaches about heme fate in Trypanosoma cruzi. Biochem. Biophys. Res. Commun. 449, 216–221

20. Hanna, D. A., Harvey, R. M., Martinez-Guzman, O., Yuan, X., Chandrasekharan, B., Raju, G., Outten, F. W., Hamza, I., and Reddi, A. R. (2016) Heme dynamics and trafficking factors revealed by genetically encoded fluorescent heme sensors. Proc. Natl. Acad. Sci. 113, 7539–44

21. Yuan, X., Rietzschel, N., Kwon, H., Walter Nuno, A. B., Hanna, D. A., Phillips, J. D., Raven, E. L., Reddi, A. R., and Hamza, I. (2016) Regulation of intracellular heme trafficking revealed by subcellular reporters. Proc. Natl. Acad. Sci. U. S. A. 113, E5144–52

22. Atamna, H., Brahmbhatt, M., Atamna, W., Shanower, G. A., and Dhahbi, J. M. (2015) ApoHRP-based assay to measure intracellular regulatory heme. Metallomics. 7, 309–321

23. Merli, M. L., Cirulli, B. A., Menendez-Bravo, S. M., and Cricco, J. A. (2017) Heme A synthesis and CcO activity are essential for Trypanosoma cruzi infectivity and replication. Biochem. J. 474, 2315–2332

24. Buchensky, C., Almirón, P., Mantilla, B. S., Silber, A. M., and Cricco, J. A. (2010) The Trypanosoma cruzi proteins TcCox10 and TcCox15 catalyze the formation of heme A in the yeast Saccharomyces cerevisiae. FEMS Microbiol. Lett. 312, 133–141

25. Sambrook, J., and Russell, D. W. (2001) Molecular Cloning: A Laboratory Manual, Molecular Cloning: A Laboratory Manual, Cold Spring Harbor Laboratory Press

26. Bookout, A. L., Cummins, C. L., Mangelsdorf, D. J., Pesola, J. M., and Kramer, M. F. (2006) High-throughput real-time quantitative reverse transcription PCR. Curr. Protoc. Mol. Biol. Chapter 15, Unit 15.8

27. Schindelin, J., Arganda-Carreras, I., Frise, E., Kaynig, V., Longair, M., Pietzsch, T., Preibisch, S., Rueden, C., Saalfeld, S., Schmid, B., Tinevez, J.-Y., White, D. J., Hartenstein, V., Eliceiri, K., Tomancak, P., and Cardona, A. (2012) Fiji: an open-source platform for biological-image analysis. Nat Meth. 9, 676–682

28. Berry, E. a, and Trumpower, B. L. (1987) Simultaneous determination of hemes a, b, and c from pyridine hemochrome spectra. Anal. Biochem. 161, 1–15

