## supporting figures 1 and 2 for "*Trypanosoma cruzi* senses intracellular heme levels and regulates heme homeostasis by modulating *Tc*HTE protein expression"

Running Tittle: *Trypanosoma cruzi* regulates heme transport *via Tc*HTE

[1]Present address: Department of Immunology and Infectious Diseases, Harvard T.H. Chan School of Public Health, 655 Huntington Avenue, FXB 217, Boston, MA 02115, USA.
[2]Present address: Department of Microbiology and Cell Science, Genetics Institute, Institute of Food and Agricultural Sciences, University of Florida, 2033 Mowry Road, PO Box 103610, Gainesville, FL 32610-3610, USA.

[3]to whom correspondence should be address: Julia A. Cricco, Instituto de Biología Molecular y Celular de Rosario (IBR), Consejo Nacional de Investigaciones Científicas y Técnicas (CONICET) – Facultad de Ciencias Bioquímicas y Farmacéuticas, Universidad Nacional de Rosario (UNR). Suipacha 531, S2002LRK, Rosario, Argentina. Tel.: +54-341-4350661 ext 133;

**Keywords:** *Trypanosoma cruzi*, Chagas disease, heme, heme transport, heme responsive gene, parasites.

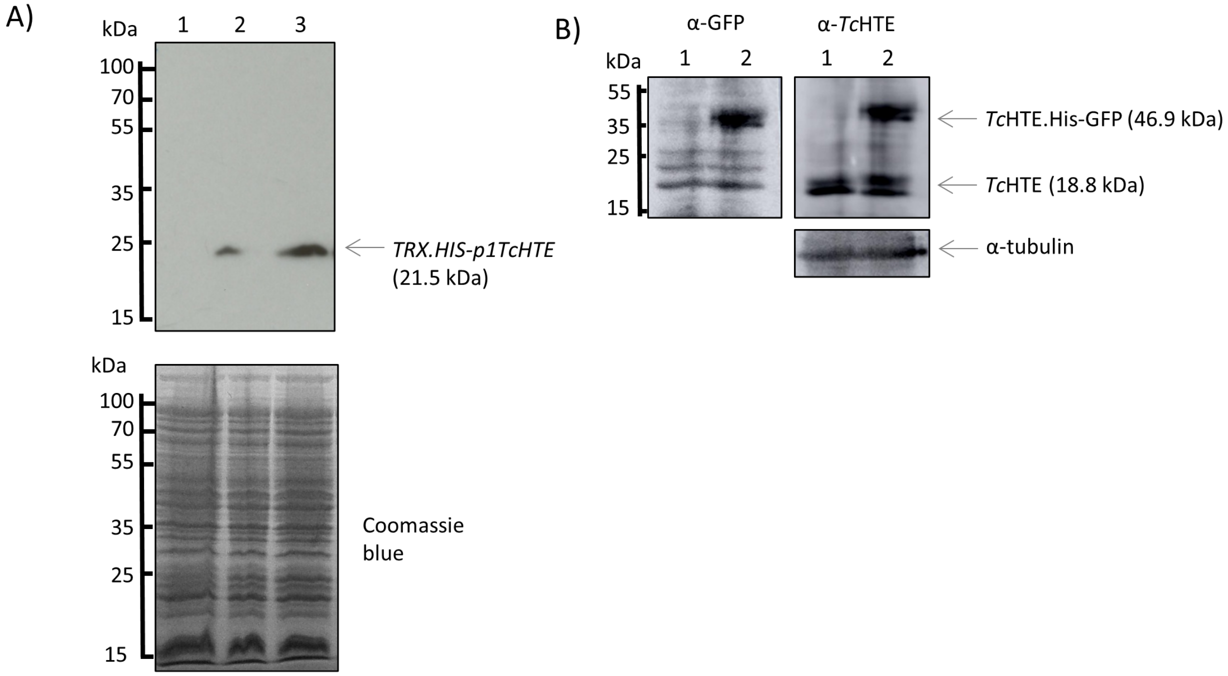


**Supporting Figure S1. A.** Extract of *E. coli* BL21(DE3)pLysS cells transformed with p*TRX.HIS-p1TcHTE* were induced with 0, 10 and 100 μM IPTG (lane 1, 2 and 3 respectively) and analyzed by Western blot using anti-*Tc*HTE immune serum (1/5000) (upper panel). The same extracts were analyzed by SDS-PAGE and stained with Coomassie blue (lower panel). **B.** Parasites were cultured in LIT-10% FBS without hemin for 48 h. WT epimastigotes (lane 1) and epimastigotes over-expressing the recombinant protein *Tc*HTE.His-GFP (lane 2) were analyzed by Western blot using anti-*Tc*HTE immune serum (1/5000) (right panel). The same membrane was striped and probed with anti-GFP antibodies (1/1000) (Santa Cruz Biotechnology) (left panel). Anti-α-tubulin was used as a loading control (lower panel). 10 x $10^6$ parasites were loaded per lane.

**Supporting Information S2. Phylogenetic analysis of HRG proteins**.

**Phylogenetic tree**. Selected sequences of putative orthologs from kinetoplastids, vertebrates and invertebrates were aligned with ClustalW (1). Transmembrane domain (TMD) predictions were performed with TMHMM Server v. 2.0 (2, 3), presenting the majority of the sequences four TMD. A cladogram was built with the sequences using Phylogeny.fr (4, 5).

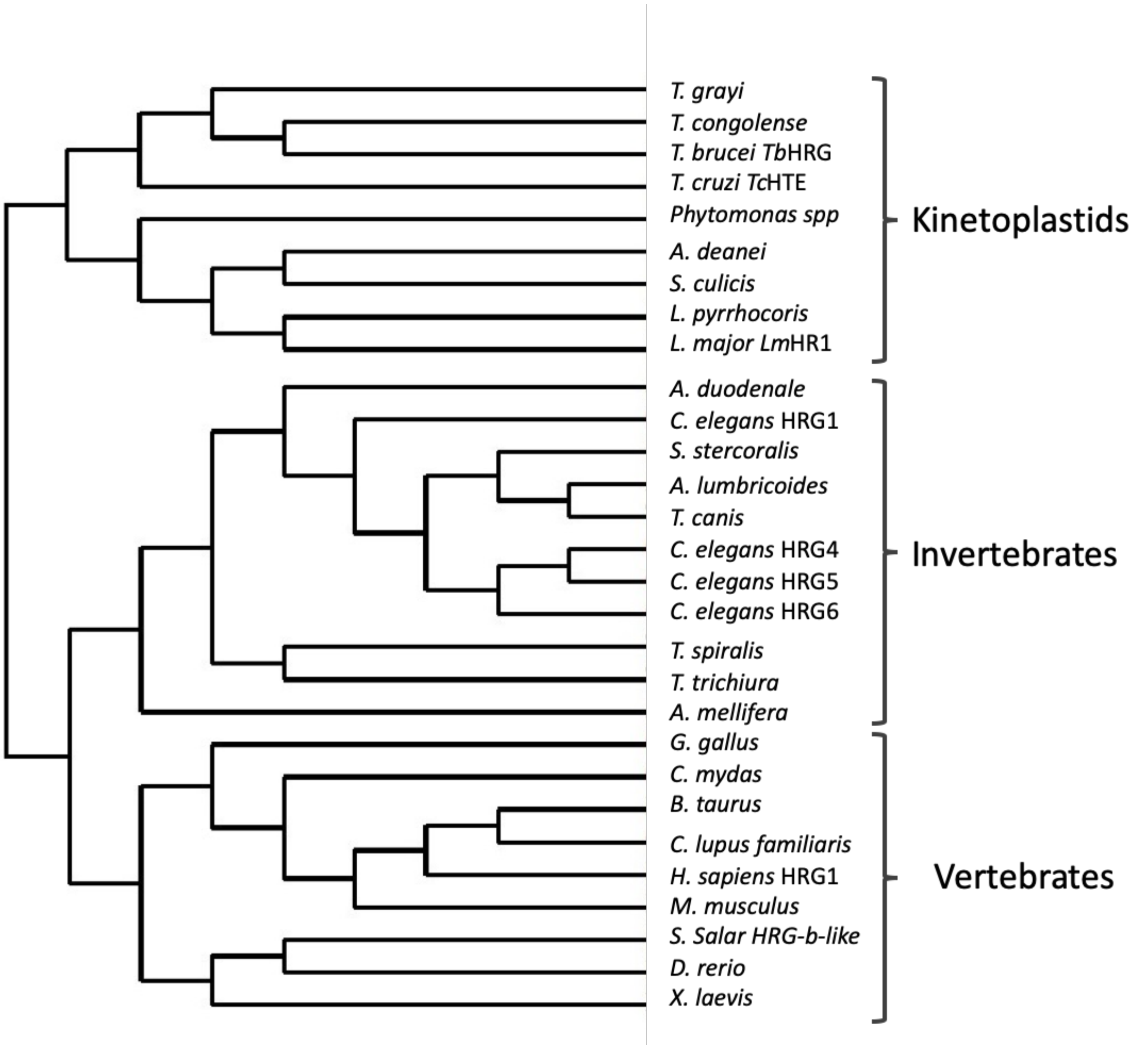


**Supporting Figure S2**. Cladogram of HRG proteins. Accession numbers: *T. grayi* GenBank Accession Number XP_009310445.1; *T. congolense* UniProt # G0USJ1; *T. brucei Tb*HRG UniProt # Q57YK2; *T. cruzi Tc*HTE UniProt # Q4DHZ7; *Phytomonas spp* UniProt # W6KM41; *A. deanei* UniProt # S9VBR5; *S. culicis* UniProt # S9VQT2; *L. pyrrhocoris* UniProt # A0A0N0DV06; *L. major Lm*HR1 UniProt # Q4QAA7; *A. duodenale* UniProt # A0A0C2G354; *C. elegans* HRG1 UniProt #

Q21642; *S. stercoralis* UniProt # A0A0K0EHS5*; A. lumbricoides* UniProt # A0A0M3I5Q9; *T. canis* UniProt # A0A183UJV1; *C. elegans* HRG-4 UniProt # Q20106; *C. elegans* HRG-5 UniProt # Q7YTM8; *C. elegans* HRG-6 UniProt # Q7YTM7; *T. spiralis* UniProt # A0A0V1BW58; *T. trichiura* UniProt # A0A077Z0X3; A. *mellifera* UniProt # A0A088AGX7; G. *gallus* UniProt # Q5ZHU0; *C. mydas* UniProt # M7AQQ6; *B. taurus* UniProt # E1BKJ0; *C. lupus familiaris* UniProt # E2RRB9; *H. sapiens* HRG1 UniProt # Q6P1K1; *M. musculus* UniProt # Q9D8M3; *S. Salar HRG-b-like* UniProt # A0A1S3MGK5; *D. rerio* UniProt # Q6ZM28; *X. Laevis* UniProt # A0A1L8HA09. All sequence IDs are UniProt accession numbers, except for *T. grayi* ID (GenBank™ accession number).

**References**


1. Larkin, M. a., Blackshields, G., Brown, N. P., Chenna, R., Mcgettigan, P. a., McWilliam, H., Valentin, F., Wallace, I. M., Wilm, a., Lopez, R., Thompson, J. D., Gibson, T. J., and Higgins, D. G. (2007) Clustal W and Clustal X version 2.0. *Bioinformatics*. **23**, 2947–2948

2. Sonnhammer, E. L., von Heijne, G., and Krogh, a (1998) A hidden Markov model for predicting transmembrane helices in protein sequences. *Proc. Int. Conf. Intell. Syst. Mol. Biol.* **6**, 175–182

3. Krogh, a, Larsson, B., von Heijne, G., and Sonnhammer, E. L. (2001) Predicting transmembrane protein topology with a hidden Markov model: application to complete genomes. *J. Mol. Biol.* **305**, 567–580

4. Dereeper, A., Audic, S., Claverie, J. M., and Blanc, G. (2010) BLAST-EXPLORER helps you building datasets for phylogenetic analysis. *BMC Evol. Biol.* **10**, 8–13

5. Dereeper, A., Guignon, V., Blanc, G., Audic, S., Buffet, S., Chevenet, F., Dufayard, J.-F., Guindon, S., Lefort, V., Lescot, M., Claverie, J.-M., and Gascuel, O. (2008) Phylogeny.fr: robust phylogenetic analysis for the non-specialist. *Nucleic Acids Res.* **36**, W465-9
